# Repeated phenotypic selection for cuticular blackness of armyworm larvae decreased stress resistance

**DOI:** 10.1101/598102

**Authors:** Takashi Matsumura, Hikaru Taya, Hitoshi Matsumoto, Yoichi Hayakawa

## Abstract

Armyworm *Mythimna separata* larvae show changes in cuticle darkening depending on population densities and are roughly categorized into two phenotypes, a pale brown solitary type and black-colored gregarious type. Although the color difference in both larval types is apparent, it remains ambiguous whether any change in physiological traits accompanies the cuticle darkening. To answer this query, we repeated genetic selection of the blackness phenotype over one hundred generations in our laboratory colony and produced a black-colored (BL) strain. Comparison between non-selected control (CTL) and BL strains revealed an increased fecundity and adult life span in the BL strain compared with the CTL strain. In contrast, BL strain larvae were found to be significantly more sensitive to heat stress than CTL strain larvae. Hemolymph reactive oxygen species (ROS) levels were higher in the BL strain than in the CTL strain irrespective of stress. Antioxidant activities of the hemolymph were not significantly different between the two strains under non-stress condition, but the activities increased to higher levels in the CTL strain than those in the BL strain after heat stress. Activities and gene expression levels of antioxidant enzymes such as catalase and superoxide dismutase (SOD) in the fat body were significantly higher in CTL strain larvae than in BL strain larvae after heat treatment. Analysis of heat stress tolerance of F1 hybrids of CTL and BL strain adults showed that phenotype of stress tolerance was inherited maternally. These results indicate a trade-off between reproductive activity and stress resistance during repeated genetic selection.

**Summary statement:** Discrete cuticle color change from whitish to blackish, which was created by repeating the reciprocal crossing of selected dark-colored individuals, increased fecundity but lowered stress tolerance in the armyworm.

## INTRODUCTION

It is known that many insect species display a phase polyphenism with obvious morphological changes that are sometimes accompanied by several metabolic and behavioral alterations (Pener and Simpson, 2009; Simpson et al., 2011). In *Locusta migratoria*, lipid metabolic profiles display significant differences between solitary and gregarious phases (Wu et al., 2012). Furthermore, in *Shistocerca gregaria*, the behavior of solitary phase individuals shifts rapidly into the gregarious state upon encountering crowded conditions with a behavioral tendency to aggregate rather than avoid one another as in the solitary state (Bazazi et al., 2008; Guttal et al., 2012).

The armyworm *Mythimna separata* shows a density-dependent polyphenism: two larval phenotypes depending on rearing population densities, a whitish solitary type and blackish gregarious type (Iwao, 1967; Ogura, 1975). It has been demonstrated that the density-dependent coloration is regulated by a specific hormone, melanization and reddish coloration hormone (MRCH) (Matsumoto et al., 1986; Suzuki et al., 1976). This color change is thought to an advantage for the insects because the black pigment allows them to absorb solar radiation, thus raising their internal temperature, and increasing their activity (Ogura 1976). In fact, it has been reported that the blackish larvae are more active, less palatable, and develop more rapidly than the whitish larvae. Such phase changes are thought to contribute toward enabling animals to rapidly adapt to environmental alterations.

In contrast with phase change-derived advantages, we have limited knowledge about the negative impacts of polyphenism in other insect species as well as the armyworm. To answer this question, we developed a blackish color (BL) strain by repeating genetic selection of the dark-colored larval phenotype under a crowded condition and compared the heat stress tolerance of this strain larvae with that of non-selected control (CTL) strain. BL strain larvae showed significantly lower resistance against heat stress compared with CTL strain larvae. A series of physiological and biochemical analyses revealed significant differences in regulatory efficiencies of redox states. In this study, we sought to reveal the mechanism underlying the phase change-derived alteration in stress tolerance of the armyworm larvae through analyses of regulatory factors of hemolymph ROS levels using larvae of both phenotype strains.

## MATERIALS AND METHODS

### Animals and heat treatments

Armyworms *Mythimna separata* (formerly called *Pseudaletia separata*) were obtained from a culture reared on an artificial diet at 25±1 °C under a photoperiod of a 16L:8D photoperiod. The diet was a kidney bean-wheat mixture supplemented with vitamins. Last (6th) instar larvae at one day after molting were used for all experiments (Matsumura et al., 2018).

In order to generate the blackish color strain, the selection of larvae with dark colored cuticles was repeated over 100 times. More than 500 selected larvae were reared in one plastic container (230 mm long, 170 mm wide, 45 mm deep) and the adults were mated with each other. In contrast, about 50 unselected larvae were raised in the same type of container and used as CTL strain larvae. Test larvae (day 1 of last instar) were exposed to heat stress conditions by placing individuals into a plastic tube (95 mm × 22 mm i.d.) and dipping it in a water bath (SDM-B, Taitec Corp. Japan) set at 38-44 °C. Controls were exposed to 25°C.

### Preparations of plasma, and fat body

Hemolymph (100 μl) was collected from three larvae into an ice-cold microcentrifuge tube containing 10 μl phosphate-buffered saline (PBS: 8 mM Na_2_ HPO_4_, 1.5 mM KH_2_ PO_4_, 137 mM NaCl, 2.7 mM KCl, pH7.2) with 0.05% N-phenylthiourea and centrifuged at 300 g for 3 min at 4 °C. Supernatant and pellet were used as a plasma and hemocyte preparations, respectively (Hayakawa, 1994). The fat body was isolated and rinsed with PBS. The tissues were blotted with filter paper and weighed before measuring ROS concentrations or enzyme activities.

### Measurements of ROS concentrations and antioxidant activities

ROS concentrations and antioxidative activities of tissue samples were measured, using an OxiSelect™In Vitro ROS/RNS Assay kit (Cell Biolabs, Inc., USA) (Cohen et al., 2016) and 2,2’-azino-bis(3-ethylbenzothiazoline-6-sulfonic acid) (ABTS) (Rubio et al., 2016), respectively, according to the procedures previously reported.

### Measurements of antioxidant enzyme activities

Catalase and SOD activities in the fat body were assayed according to the procedures previously described with minor changes. Briefly, fat bodies dissected from test larvae were sonicated in 50 mM phosphate buffer (pH 7.0) containing 1 mM EDTA by an ARTEK Sonic Dismembrator (10 pulses at 50W), and the supernatant was centrifuged at 5,000 *g* for 10 min at 4°C before it was used as the crude enzyme preparation. The catalase activity of each sample was measured as follows (Johansson and Borg, 1988). After 10 μl of the enzyme sample was mixed with 125 μl assay buffer (100 mM potassium phosphate buffer, pH 7.0), 50 μl methanol, and 10 μl hydrogen peroxide (27%), the mixture was incubated for 10 min under continuous shaking at 20°C. The enzyme reaction was terminated by adding 39 μl 10 M potassium hydroxide and 74.3 μl Purpald (46 mM in 0.5 M HCl), and incubated for 10 min under continuous shaking at 20°C. At 5 min after adding 17 μl 192 mM potassium periodate to oxidize the Purpald-formaldehyde complex, absorbance was measured at 450 nm. One unit of catalase activity catalyzed the degradation of 1 μmol of H_2_ O_2_ per min.

SOD activity was quantified as follows (Peskin and Winterbourn, 2000). Ten μl of the enzyme sample was mixed with 250 μl 50 mM Tris-HCl buffer (pH 8.0) containing 50 mM diethylene-triamine-penta-acetic acid, 50 mM hypoxanthine, 52 μM WST-1, and 240 mU/ml xanthine oxidase (Sigma-Aldrich Co., USA). After incubation at 37 °C for 20 min, absorbance was measured at 450 nm. One unit of SOD activity was defined as the amount required for inhibiting the tetrazolium reduction by 50% per min.

### Quantitative RT-PCR (qPCR)

Total RNA was extracted from each tissue using TRIzol (Gibco-BRL, USA) according to the manufacturer’s protocol. First-strand cDNA was synthesized with an oligo(dT)_12-18_ primer using ReverTra Ace RT-PCR kit (Toyobo) according to the manufacturer’s protocol. qPCR analysis of each gene was carried out using a Light-Cycler 1.3 instrument (Roche Applied Science) under the following conditions: 45 cycles at 95°C for 10 sec, 55°C for 10 sec, and 72°C for 20 sec (Tsuzuki et al., 2014; Tsuzuki et al., 2012). PCR specificity was confirmed by sequencing the PCR products and melting curve analysis at each data point. All samples were analyzed in duplicate, and assay variation was typically within 10%. Data were normalized to the expression level of *rp49* determined in duplicate by reference to a serial dilution calibration curve (Bustin et al., 2009). Primer pairs of target genes indicated in Results were: *rp49*, TGACAAACTCAAGCGTAACTGGC and TTGCGGAAACCATTGGGCAG; *catalase*, AAAGGAGCTGGAGCTTT and CTTCTGAGTATGGATAAA; *SOD1*, TGGCTGCATGTCGTC and AGATCATCGGCATCGG; *SOD2*, CCACATCAACCACACC and CGTTCTTGTACTGCAG.

### Statistical analysis

Statistical analyses were carried out using Tukey’s HSD tests, unless otherwise stated. The Shapiro-Wilk test confirmed that data sets do not deviate from normality. P values of less than 0.05 were regarded as statistically significant. All statistical analyses were performed using JMP 9.0.2 (SAS Institute).

## RESULTS

### Physiological traits of CTL and BL strains

We generated BL strain larvae by repeated genetic selection of the blackish colored phenotype, as shown in Fig. 1A. BL strain adults showed higher fecundity than the CTL strain: BL strain females laid approximately 3 times more eggs as the CTL strain did (Fig. 1B). Consistent with the enhanced fecundity of the BL strain, the adult life span was also longer in the BL strain than in the CTL strain when equal numbers of males and females were reared together in a cage (Fig. 1C).

**Fig. 1.**
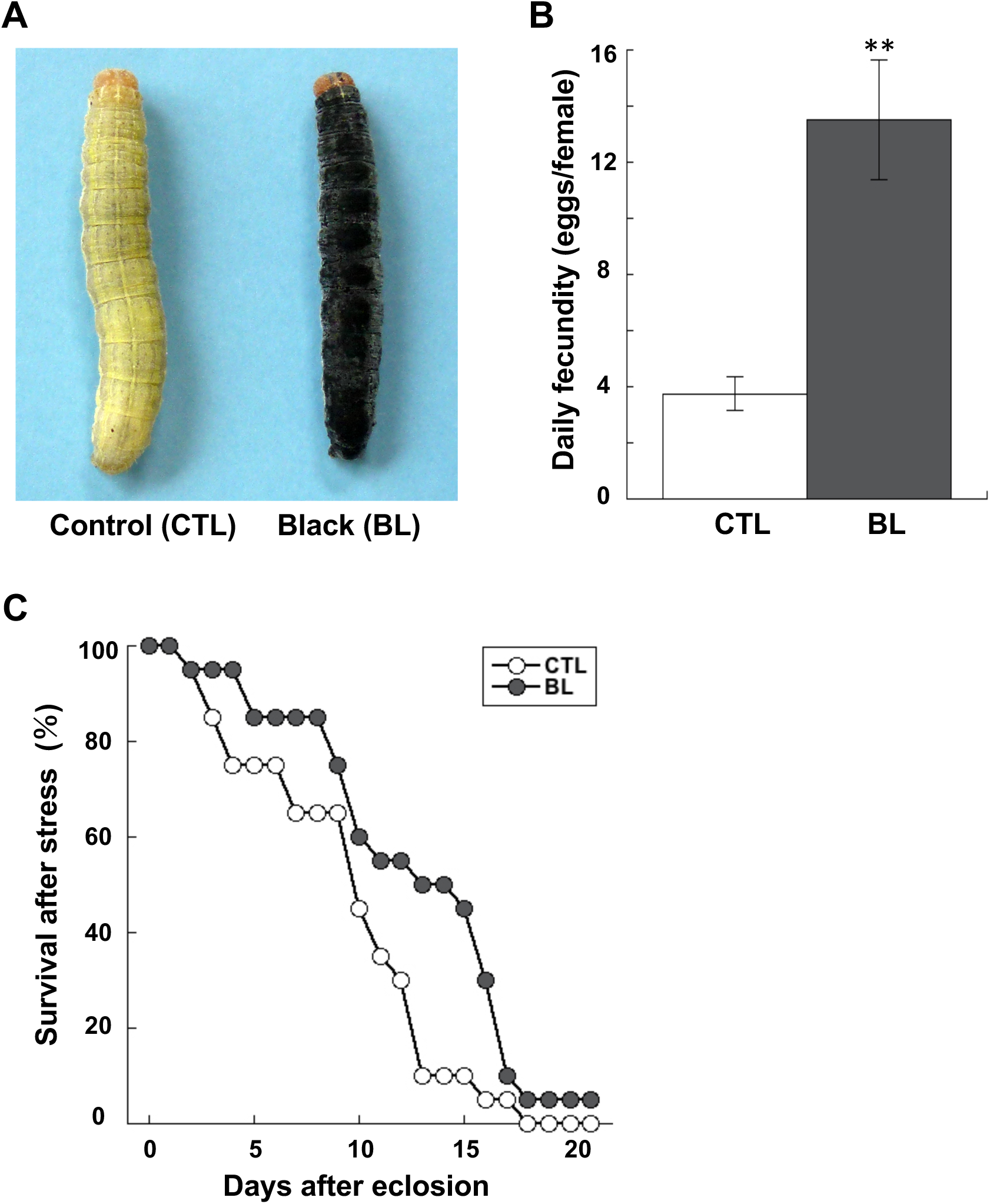
Phenotypic differences in unselected control whitish (CTL) and selected blackish (BL) strains. (A) Images of CTL and BL strain last instar larvae. (B) The average number of eggs laid by one mated female a day (data are means ± SEM; n=25). **P<0.01 vs. CTL larvae. (C) Increased life spans of BL strain adults. Five newly eclosed male and female adults were reared together in a cage (35.5 cm × 36.5 cm × 25.5 cm) containing 10% honey solution. The number of dead moths was counted every day (data are means ± SEM; n=10). Significant difference of both slopes was determined using the log-rank Mantel Cox Test (P<0.05).

Comparison of heat stress tolerance in both strains showed that BL strain larvae were more sensitive to heat stress at 43.5°C for 1 h than CTL strains (Fig. 2A). Measurement of survivals of both strains of larvae after heat treatments at 43.5°C for various periods showed that 4 hour’s heat stress caused 100% mortality in the BL strain larvae but approximately 60% mortality in CTL larvae (Fig. 2B). When both strains of larvae were exposed to 1 hour’s heat stress at various temperatures, median lethal temperatures (LT50) of CTL and BL strains were 42.8 °C and 43.9 °C, respectively (Fig. 2C).

**Fig. 2.**
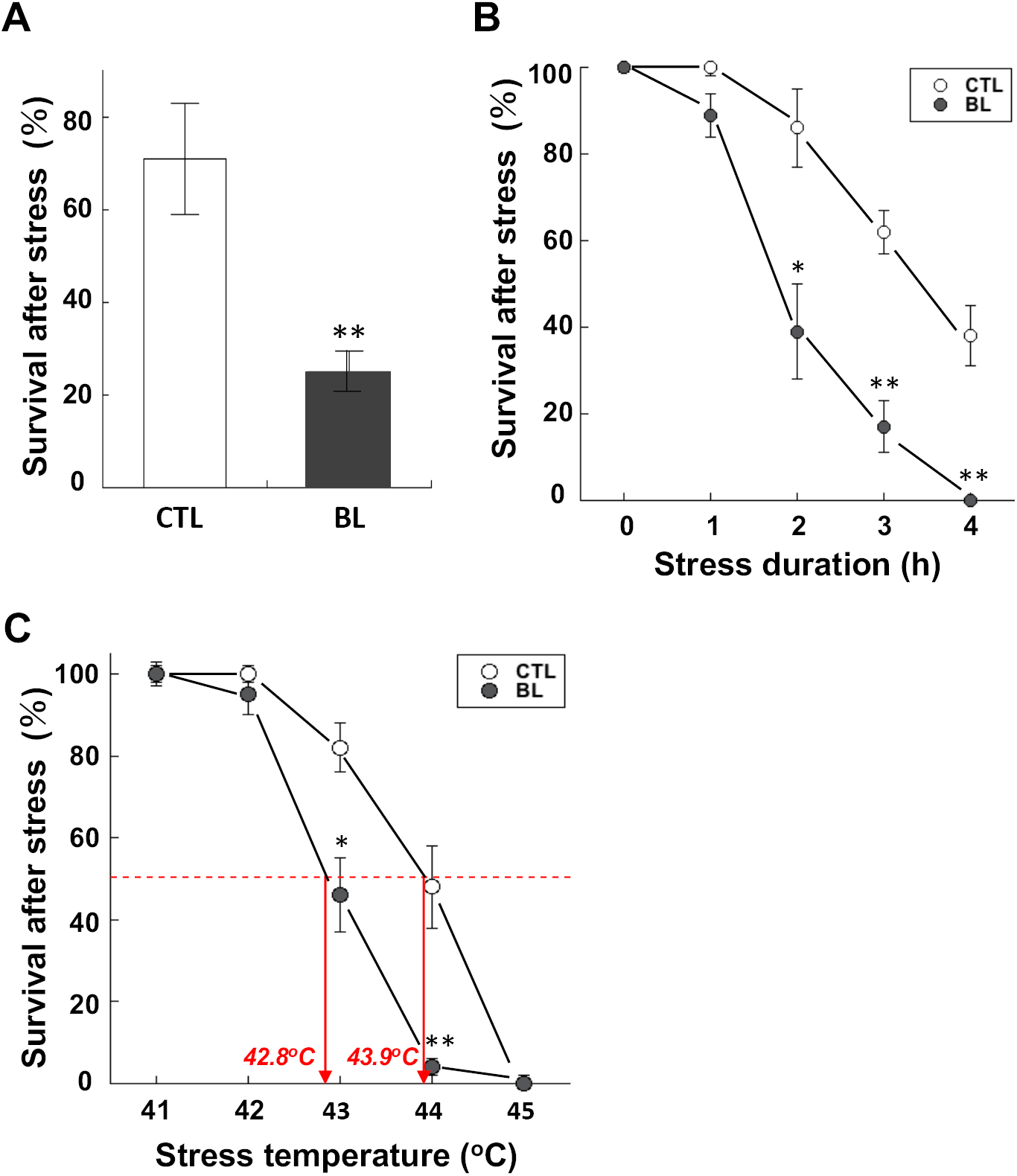
Heat stress tolerance of CTL and BL strain larvae. (A) Survival of CTL and BL strain larvae 2days after heat stress at 43.5°C for 1 h (data are means ± SEM; n=20). **P<0.01 vs. CTL. (B) Survival of CTL and BL strain larvae 1 days after heat stress at 43.5°C for 0-4 h (data are means ± SEM; n=6). *P<0.05, **P<0.01 vs. CTL. (C) Survival of CTL and BL strain larvae 2 days after heat stress at 41-44°C for 1 h (data are means ± SEM; n=8). *P<0.05, **P<0.01 vs. CTL.

### Hemolymph ROS and antioxidant activity changes after heat stress

ROS concentrations in the hemolymph of CTL and BL strain larvae were measured before and after incubation at 43.5°C for 1 h. Hemolymph ROS concentrations were found to be much higher in BL strain larvae than in CTL strain larvae irrespective of stress, and kept increasing in both strains of larvae after heat stress (Fig. 3A).

**Fig. 3.**
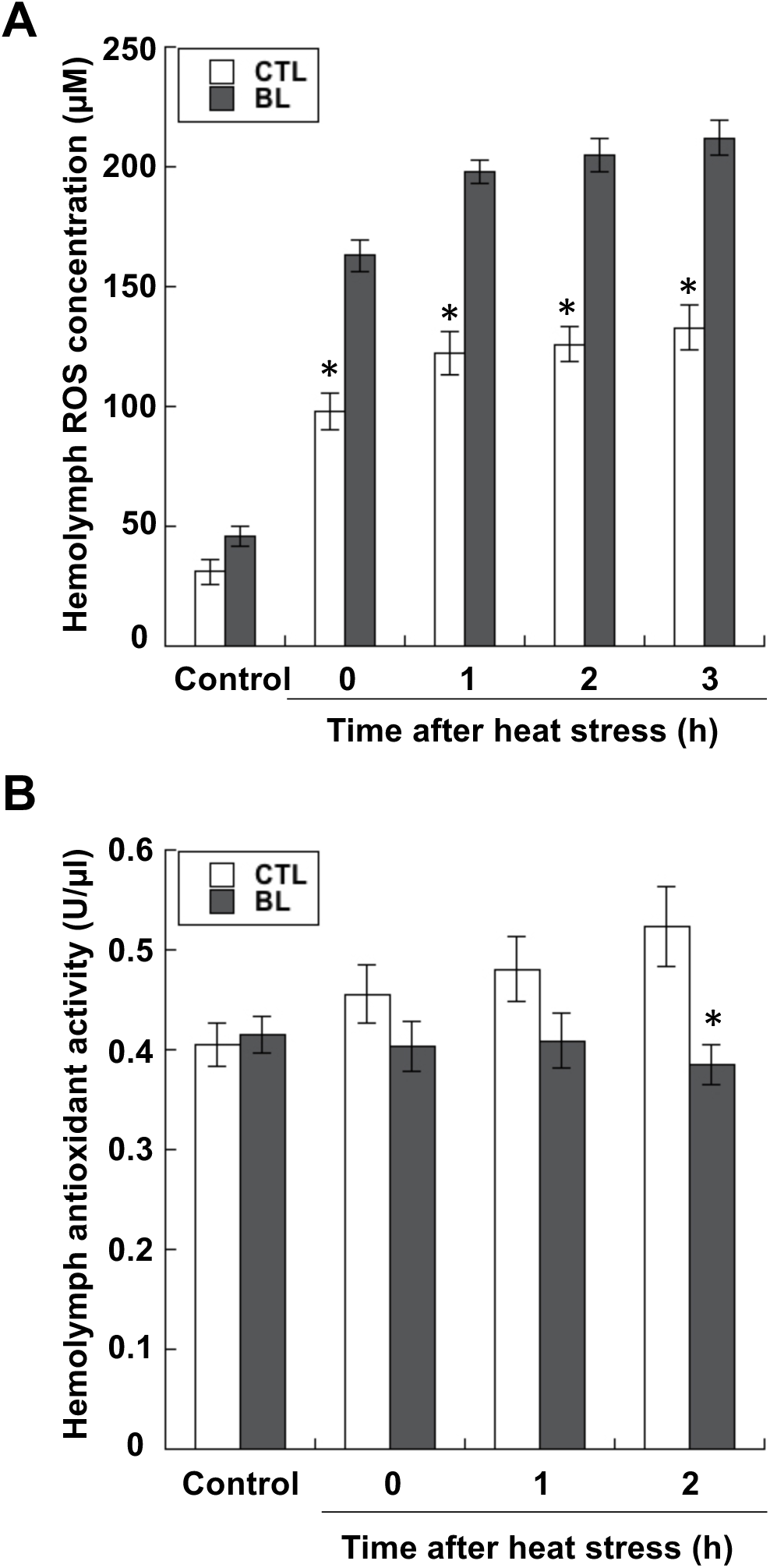
ROS concentrations and antioxidant activities in the hemolymph of CTL and BL strain larvae. (A) ROS concentrations in the hemolymph of both strain larvae before (Control) and after heat stress at 43.5°C for 1 h (data are means ± SEM; n=6). *P<0.05 vs. Control. (B) Antioxidant activities in the hemolymph of both strain larvae before (Control) and after heat stress at 43.5°C for 1 h (data are means ± SEM; n=6). *P<0.05 vs. Control.

Antioxidant activities in the hemolymph of both strains were not significantly different before heat treatment (Fig. 3B). After heat stress, antioxidant activities increased in the hemolymph of CTL strain larvae, but they did not change in the BL strain.

### Antioxidant enzyme activities and gene expression after heat stress

SOD activities in the fat body of CTL strain larvae were significantly higher than in the BL strain both before and after heat stress. Furthermore, the increase in SOD activities after heat treatment was observed in the CTL larval fat body but not in the BL larval fat body (Fig. 4A). Catalase activities in the fat body were not significantly different between both strain larvae before heat treatment, while, after heat stress, catalase activities increased in CTL larvae but not in the BL strain (Fig. 4B).

**Fig. 4.**
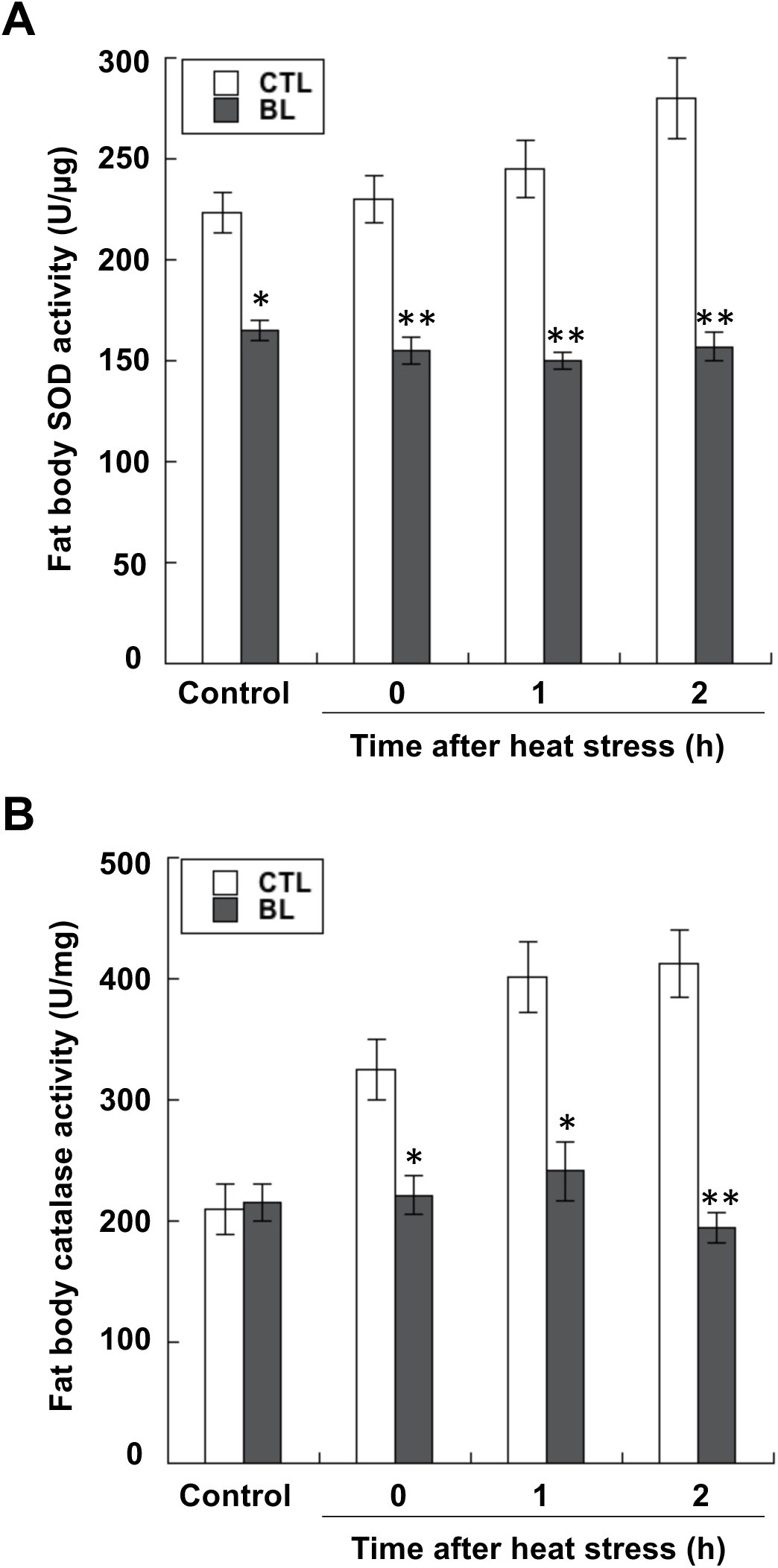
Fat body SOD and catalase activities of CTL and BL strain larvae. (A) Fat body SOD activities of both strain larvae before (Control) and after heat stress at 43.5°C for 1 h (data are means ± SEM; n=5). *P<0.05, **P<0.01 vs. Control. (B) Fat body catalase activities of both strain larvae before (Control) and after heat stress at 43.5°C for 1 h (data are means ± SEM; n=5). *P<0.05, **P<0.01 vs. Control.

*SOD1* expression levels in the fat body were much higher in CTL larvae than in BL larvae before and after heat treatment (Fig. 5A). Although *SOD1* expression levels in BL larval fat bodies were not changed after heat stress, the expression in CTL larval fat bodies increased after heat stress. Fat body *SOD2* expression levels were also much higher in CTL strain larvae than in BL larvae both before and after heat treatment (Fig. 5B). The expression levels in both strains were not changed after heat stress.

**Fig. 5.**
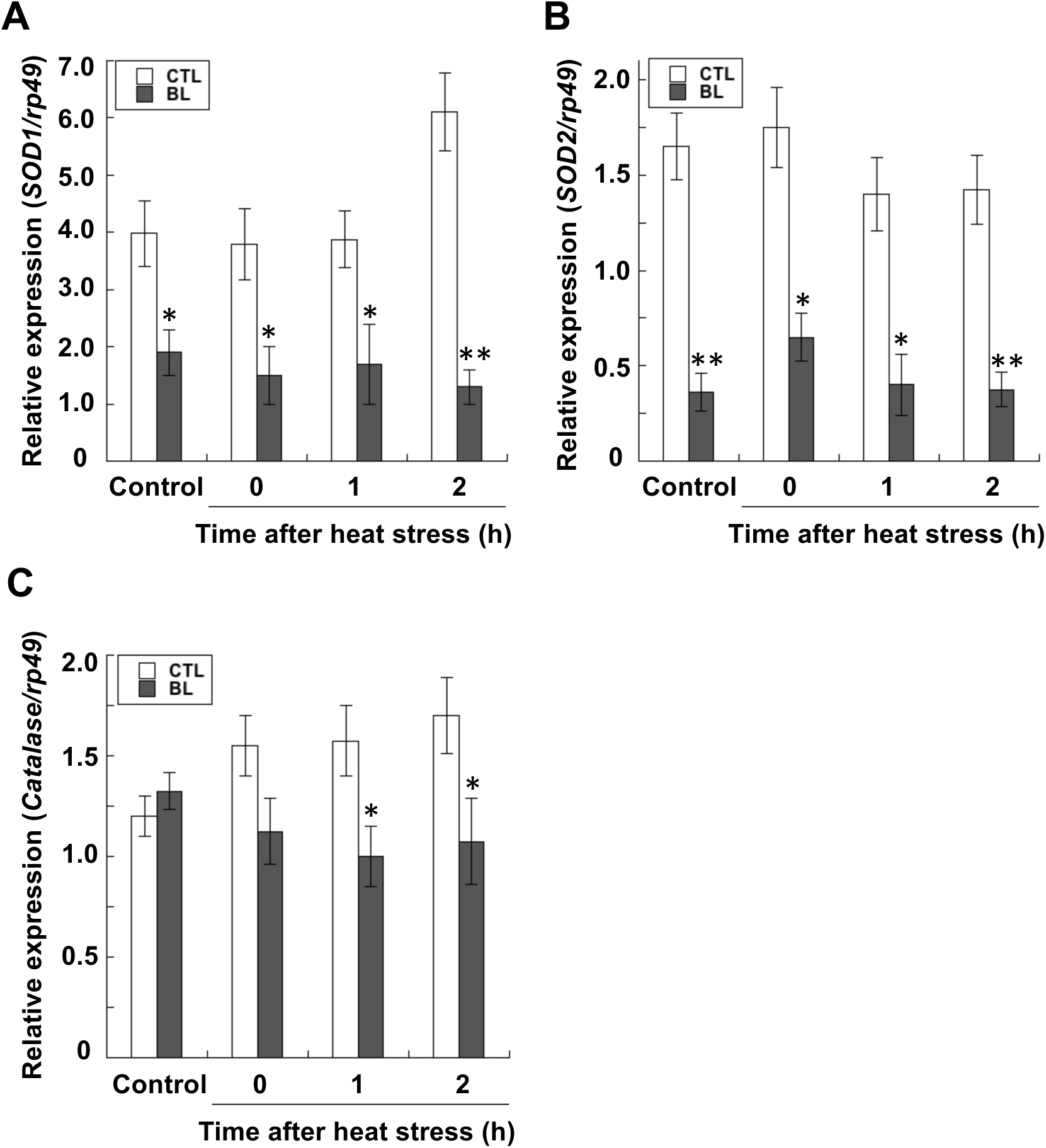
*SOD 1, SOD2*, and *catalase* expression levels in the fat body of CTL and BL strain larvae. (A) *SOD1* expression levels in the fat body of both strain larvae before (Control) and after heat stress at 43.5°C for 1 h (data are means ± SEM; n=7). *P<0.05, **P<0.01 vs. Control. (B) *SOD2* expression levels in the fat body of both strain larvae before (Control) and after heat stress at 43.5°C for 1 h (data are means ± SEM; n=7). *P<0.05, **P<0.01 vs. Control. (C) *Catalase* expression levels in the fat body of both strain larvae before (Control) and after heat stress at 43.5°C for 1 h (data are means ± SEM; n=7). *P<0.05 vs. Control.

Fat body *catalase* expression levels were not significantly different between CTL and BL strains before heat treatment (Fig. 5C). After heat stress, *catalase* expression increased in CTL strain larvae but did not in BL larvae.

### Stress tolerance of F1 hybrids of CTL and BL strain adults

Reciprocal crosses were made between CTL and BL strains, and F1 larvae were exposed to heat stress to compare their stress tolerance. The survivals of F1 crossbred (CTL female x BL male: CF-F1) larvae were much higher than those of reciprocal F1 crossbred larvae (Fig. 6A), indicating that the stress tolerance is likely inherited maternally.

**Fig. 6.**
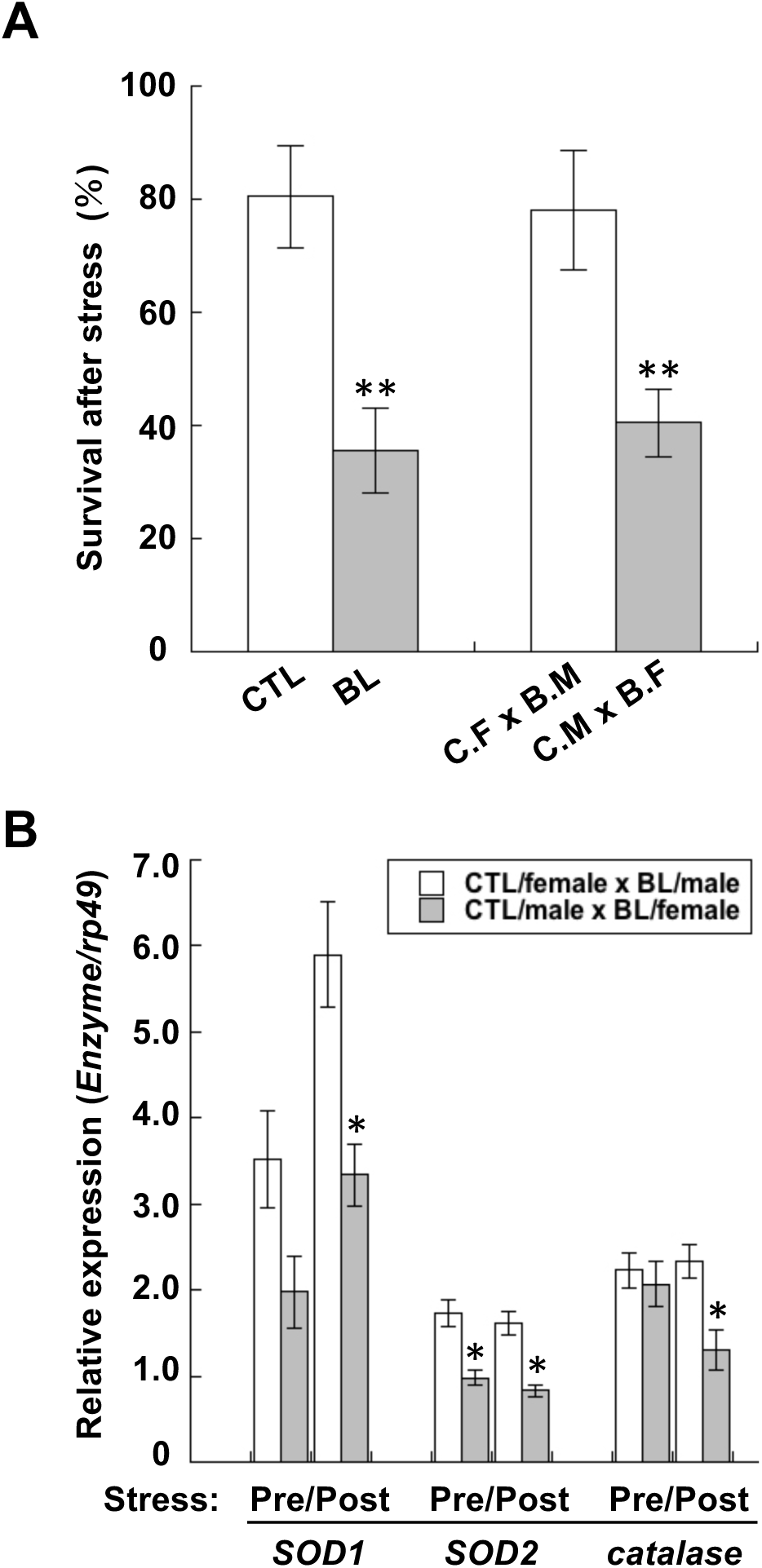
Stress tolerance and antioxidant enzyme gene expression levels of CTL strain larvae, BL strain larvae, and their F1 hybrid larvae. (A) Survival of CTL strain larvae, BL strain larvae, and their F1 hybrid larvae after heat stress at 43.5°C for 1 h (data are means ± SEM; n=6). **P<0.01 vs. Control. C.F. x B.M.: CTL/female x BL/male; C.M. x B.F.: CTL/male x BL/female. (B) *SOD1, SOD2*, and *catalase* expression levels in the fat body of larvae of F1 hybrid between before (Pre) and after (Post) heat stress at 43.5°C for 1 h (data are means ± SEM; n=5). *P<0.05 vs. CTL/female x BL/male.

Measurements of antioxidant enzyme gene expression before and after heat stress showed that the expression patterns of the three genes in CF-F1 was more or less similar to those of the CTL strain, although the stress-induced increase in *SOD1* expression in CF-1 was less than that in CTL strain (Fig. 6B).

## DISCUSSION

The cuticle color of armyworm larvae is an inheritable trait, which allowed us to generate strain BL in our laboratory. We created this strain by repeating their reciprocal crossing of selected dark-colored individuals. We found that the BL strain has higher reproducibility compared with the CTL strain: the average number of eggs laid by one BL female was approximately three times that laid by one CTL female. This would be at least partly due to the longer life span of the BT strain compared to the CTL strain but mostly due to higher egg production rates by BR strain females compared with CTL females because the difference in life spans between the strains was not much.

Comparison of stress tolerance in both strain larvae showed higher sensitivities of BL strain larvae to heat stress compared with CTL strain larvae. The TL50 of BL strain larvae was 42.8°C, which is one degree lower than the TL 50 (43.9°C) of the CTL strain. Measurement of hemolymph ROS concentrations demonstrated that BL strain larvae possessed higher ROS than CTL larvae. It is reasonable to assume that higher concentrations of ROS in the BL strain larvae are due to higher production and lower degradation of ROS. We speculate that the higher production of ROS in the BL strain would be attributed chiefly to higher production of melanin in BL strain larvae compared with CTL strain larvae. In fact, our preliminary experiments showed much higher gene expression levels of both tyrosine hydroxylase and dopa decarboxylase, which play a major role in dopamine melanin synthesis in insects, in BL strain larvae than in CTL strain larvae. In insects, both enzymes catalyze tyrosine to dopamine, and subsequent oxidation and polymerization results in the synthesis of melanin (Ninomiya and Hayakawa, 2007; Ninomiya et al., 2008; Ninomiya et al., 2006; True et al., 1999), during which these reactions are associated with ROS production (Munoz-Munoz et al., 2009). Primary antioxidant enzymes such as SODs and catalase degrade ROS in insects (Ahmad and Pardini, 1990). We confirmed that both enzyme activities are higher in the fat body of CTL strain larvae than BL strain larvae. Differences between the activities of the two strains were magnified after heat stress, suggesting that CTL strain larvae potentially have a higher ability to adapt to changes in their environments compared with BL strain larvae. It is reasonable to assume that such a high adaptability of the CTL strain is attributable to genetic variation such as changes in a certain transcription factor expression level because we confirmed that expression levels of these enzyme genes were significantly higher in CTL strain larvae than in BL strain larvae, especially after heat stress. Our prior report showing the enhanced expression of in *SODs* and *catalase* in the armyworm larvae acclimated to thermal stress by pretreatment with mild heat stress is consistent with supporting this interpretation (Matsumura et al., 2017).

In some insect species as well as mammals, expression of *SODs* and *catalase* has been reported to be regulated by FoxO transcription factor(s) (Klotz et al., 2015; Lu et al., 2018; Sim and Denlinger, 2011). We preliminarily observed that transcriptional enhancement of FoxO and subsequent translocation into the nuclei occurred in the fat body of *Drosophila* larvae during heat stress acclimation. Therefore, it is possible that expression levels of FoxO as well as its localization would be different between CTL and BL strain larvae: FoxO would be more active in the former than the latter. Further studies aimed at revealing the upstream regulatory mechanism of the antioxidant gene expression should enhance understanding of the mechanisms underlying the difference in stress tolerance between the two strains.

We previously compared stress tolerance between two strains of the red flour beetle, *Tribolium castaneum*, with long tonic immobility (average 377 s immobility duration: L-type) and short tonic immobility (average 1.25 s immobility duration: S-type) (Kiyotake et al., 2014). Although tonic immobility (death-feigning) behavior has been demonstrated to be an effective predator defense mechanism (Miyatake et al., 2004), we found that only 6.5% of the beetles collected in the storehouse showed over 60 s duration of death-feigning, while 70% of the collected beetles showed less than 10 s duration. Comparison of stress tolerance between two strains under several stressors including low temperature, high temperature, and mechanical vibration demonstrated that L-type beetles were significantly more sensitive to all tested stresses. Moreover, we found that expression levels of *catalase* and insect cytokine *growth-blocking peptide* (Hayakawa, 1990, 1991) were significantly higher in S-type than in L-type beetles. This story shares close similarities with the present observation because, although both phenotypes, long death-feigning of the beetles and darkening cuticle color of the armyworm, had acquired the merit of defending themselves under particular conditions such as facing predators and crowding, both of them had lost stress tolerance. Furthermore, the key genes that contributed to such a phenotype alteration are similar: it is likely attributable to antioxidant enzyme gene(s). Therefore, these two stories could be considered as similar examples of trade-offs between stress tolerance and competitive abilities in these insect species.

## Abbreviation

GBP: growth-blocking peptide
ROS: reactive oxygen species
SOD: superoxide dismutase

## ACKNOWLEDGEMENTS

This work was supported by a Grant-in-Aid for Scientific Research (A) (Grant number: 16H0259) from JSPS.

## REFERENCES

Ahmad, S., Pardini, R., 1990. Mechanisms for regulating oxygen toxicity in phytophagous insects. Free Radic. Biol. Med. 8, 401–413.

Bazazi, S., Buhl, J., Hale, J.J., Anstey, M.L., Sword, G.A., Simpson, S.J., Couzin, I.D., 2008. Collective motion and cannibalism in locust migratory bands. Curr. Biol. 18, 735–739.

Cohen, S., Mehrabi, S., Yao, X., Millingen, S., Aikhionbare, F.O., 2016. Reactive oxygen species and serous epithelial ovarian adenocarcinoma. Cancer Res. J. 4, 106.

Guttal, V., Romanczuk, P., Simpson, S.J., Sword, G.A., Couzin, I.D., 2012. Cannibalism can drive the evolution of behavioural phase polyphenism in locusts. Ecol. Lett. 15, 1158–1166.

Hayakawa, Y., 1990. Juvenile hormone esterase activity repressive factor in the plasma of parasitized insect larvae. J. Biol. Chem. 265, 10813–10816.

Hayakawa, Y., 1991. Structure of a growth-blocking peptide present in parasitized insect hemolymph. J. Biol. Chem. 266, 7981–7984.

Hayakawa, Y., 1994. Cellular immunosuppressive protein in the plasma of parasitized insect larvae. J. Biol. Chem. 269, 14536–14540.

Iwao, S.i., 1967. Differences in light reactions of larvae of the armyworm, *Leucania separata* Walker, in relation to their phase status. Nature 213, 941–942.

Johansson, L.H., Borg, L.H., 1988. A spectrophotometric method for determination of catalase activity in small tissue samples. Anal. Biochem. 174, 331–336.

Kiyotake, H., Matsumoto, H., Nakayama, S., Sakai, M., Miyatake, T., Ryuda, M., Hayakawa, Y., 2014. Gain of long tonic immobility behavioral trait causes the red flour beetle to reduce anti-stress capacity. J. Insect Physiol. 60, 92–97.

Klotz, L.O., Sanchez-Ramos, C., Prieto-Arroyo, I., Urbanek, P., Steinbrenner, H., Monsalve, M., 2015. Redox regulation of FoxO transcription factors. Redox Biol. 6, 51–72.

Lu, J., Wang, Z., Cao, J., Chen, Y., Dong, Y., 2018. A novel and compact review on the role of oxidative stress in female reproduction. Reprod. Biol. Endocrinol. 16, 80.

Matsumoto, S., Isogai, A., Suzuki, A., 1986. Isolation and amino terminal sequence of melanization and reddish coloration hormone (MRCH) from the silkworm, *Bombyx mori*. Insect Biochem. 16, 775–779.

Matsumura, T., Matsumoto, H., Hayakawa, Y., 2017. Heat stress hardening of oriental armyworms is induced by a transient elevation of reactive oxygen species during sublethal stress. Arch. Insect Biochem. Physiol. 96, e21421.

Matsumura, T., Nakano, F., Matsumoto, H., Uryu, O., Hayakawa, Y., 2018. Identification of a cytokine combination that protects insects from stress. Insect Biochem. Mol. Biol. 97, 19–30.

Miyatake, T., Katayama, K., Takeda, Y., Nakashima, A., Sugita, A., Mizumoto, M., 2004. Is death-feigning adaptive? Heritable variation in fitness difference of death-feigning behaviour. Proc. R. Soc. B: Biol. Sci. 271, 2293–2296.

Munoz-Munoz, J.L., Garcia-Molina, F., Varon, R., Tudela, J., Garcia-Canovas, F., Rodriguez-Lopez, J.N., 2009. Generation of hydrogen peroxide in the melanin biosynthesis pathway. Biochim. Biophys. Acta 1794, 1017–1029.

Ninomiya, Y., Hayakawa, Y., 2007. Insect cytokine, growth-blocking peptide, is a primary regulator of melanin-synthesis enzymes in armyworm larval cuticle. FEBS J. 274, 1768–1777.

Ninomiya, Y., Kurakake, M., Oda, Y., Tsuzuki, S., Hayakawa, Y., 2008. Insect cytokine growth-blocking peptide signaling cascades regulate two separate groups of target genes. FEBS J. 275, 894–902.

Ninomiya, Y., Tanaka, K., Hayakawa, Y., 2006. Mechanisms of black and white stripe pattern formation in the cuticles of insect larvae. J. Insect Physiol. 52, 638–645.

Ogura, N., 1975. Hormonal control of larval coloration in the armyworm, *Leucania separata*. J. Insect Physiol. 21, 559–576.

Ogura, N., 1976. Role of oesophageal connectives in cuticular melanization in larvae of the armyworm, *Leucania separata* WALKER (Lepidoptera: Noctuidae). Appl. Entomol. Zool. 11, 70–74.

Pener, M.P., Simpson, S.J., 2009. Locust phase polyphenism: an update. Adv. Insect Physiol. 36, 1–272.

Peskin, A.V., Winterbourn, C.C., 2000. A microtiter plate assay for superoxide dismutase using a water-soluble tetrazolium salt (WST-1). Clin. Chim. Acta 293, 157–166.

Rubio, C.P., Hernández-Ruiz, J., Martinez-Subiela, S., Tvarijonaviciute, A., Ceron, J.J., 2016. Spectrophotometric assays for total antioxidant capacity (TAC) in dog serum: an update. BMC Vet. Res. 12, 166.

Sim, C., Denlinger, D.L., 2011. Catalase and superoxide dismutase-2 enhance survival and protect ovaries during overwintering diapause in the mosquito *Culex pipiens*. J. Insect Physiol. 57, 628–634.

Simpson, S.J., Sword, G.A., Lo, N., 2011. Polyphenism in insects. Curr. Biol. 21, R738–R749.

Suzuki, A., Matsumoto, S., Ogura, N., Isogai, A., Tamura, S., 1976. Extraction and partial purification of the hormone inducing cuticular melanization in armyworm larvae. Agric. Biol. Chem. 40, 2307–2309.

True, J.R., Edwards, K.A., Yamamoto, D., Carroll, S.B., 1999. *Drosophila* wing melanin patterns form by vein-dependent elaboration of enzymatic prepatterns. Curr. Biol. 9, 1382–1391.

Tsuzuki, S., Matsumoto, H., Furihata, S., Ryuda, M., Tanaka, H., Sung, E.J., Bird, G.S., Zhou, Y., Shears, S.B., Hayakawa, Y., 2014. Switching between humoral and cellular immune responses in *Drosophila* is guided by the cytokine GBP. Nat. Commun. 5, 4628.

Tsuzuki, S., Ochiai, M., Matsumoto, H., Kurata, S., Ohnishi, A., Hayakawa, Y., 2012. *Drosophila* growth-blocking peptide-like factor mediates acute immune reactions during infectious and non-infectious stress. Sci. Rep. 2, 210.

Wu, R., Wu, Z., Wang, X., Yang, P., Yu, D., Zhao, C., Xu, G., Kang, L., 2012. Metabolomic analysis reveals that carnitines are key regulatory metabolites in phase transition of the locusts. Proc. Nat. Acad. Sci. 109, 3259–3263.

